# A minimally-invasive method for serial cerebrospinal fluid collection and injection in rodents with high survival rates

**DOI:** 10.1101/2022.09.30.510413

**Authors:** Jingrong Regina Han, Yu Yang, Tianshu William Wu, Tao-Tao Shi, Wenlu Li, Yilong Zou

## Abstract

Cerebrospinal fluid (CSF) is a clear fluid surrounding and nourishing the brain and spinal cord. Molecular profiling of the CSF is a common diagnostic approach for central nervous system (CNS) diseases, including infectious diseases, autoimmune disorders, brain hemorrhage and traumatic brain injury, CNS tumors, and Alzheimer’s disease^1–10^. Rodent models are critical for investigating CNS disease mechanisms and therapeutics, however, both collecting CSF and injecting materials into CSF in small animals are technically challenging and often result in high rates of postoperative mortality. Here, we present an easy-to-practice and cost-effective protocol with minimum instrument requirements to access the CSF in live rodents for collection and infusion purposes. By introducing a metal needle tool bent at a unique angle and length, we could steadily reach the CSF via the foramen magnum. Compared with prior methods, this protocol requires neither the operator to discern the changes in resistance from solid tissues while puncturing the needle, nor surgical opening of the skin and muscle covering the rodent neck. Using this method, we frequently obtain 5-15 μL of CSF from mice and 70-120 μL from rats to enable diverse downstream analyses including mass spectrometry. Due to the minimal invasiveness, this procedure allows iterative CSF collection from the same animal every few days – a major improvement over prior protocols that require extensive surgical operations. Moreover, we demonstrate that this method could be used for injecting desired solutions including dyes into mouse CSF with high success rates. Our method shortens the time required for CSF collection or injection to 3-5 minutes. Notably, we could reach near 100% postoperative recovery rates in both mice and rats even with repetitive collections. Together, we establish an efficient and minimally-invasive protocol for collecting CSF and inoculating reagents into the CSF in live rodents to enable various longitudinal studies at the forefronts of CNS investigation.

## Introduction

Diagnosis of central nervous system (CNS) diseases is a continuous challenge in biomedicine. The cerebrospinal fluid (CSF) provides an important window and sample source for discovering biomarkers to read out the pathological states of the CNS. Under physiological conditions, the CSF is predominantly secreted by the choroid plexus within the ventricular interior of the brain to circulate in the ventricles and subarachnoid space of the brain and spinal cord^11^. The CSF supports CNS functions by supplying nutrients, removing waste, transduction of signals, and protecting against pathogens^11,12^. In the presence of mechanical impact, the CSF also serves as a cushion to protect the brain and spinal cord tissues from acute injury. Due to the pleiotropic functions of the CSF, compositional analysis of the CSF via multi-omics or targeted approaches is widely used to diagnose and prognose CNS disorders^13^. In addition to the diagnostic value, recent research revealed that transfering young CSF into aged mice could restore memory^14^, highlighting the therapeutic potential of CSF in treating CNS related diseases. Nevertheless, extracting CSF from small laboratory animals including mice remains a challenge due to small body sizes and limited CSF volume.

Currently, there are three typical routes for CSF collection in mice, including accessing the liquid phase through the cisterna magna, the lateral ventricle and the intrathecal route^15–17^. The cisterna magna method requires a fine and firm puncture by the operator, and is not compatible with repeated collection of CSF. Likewise, the lateral ventricle collection method requires using a thin metal needle to penetrate the brain parenchyma for stereotaxic targeting towards the ventricle space, which can be inaccurate and may cause brain injury. On the other hand, intrathecal cannulation may cause injury to the brain parenchymal tissue due to cannula insertion. Collectively, the existing CSF collection methods require both sophisticated instruments and well-trained operators, and could lead to high mortality rates or reach euthanization endpoints for animal welfare^15–17^. These limitations prevent the exploration of CNS disorders and longitudinal studies that require consecutive sampling of the CSF from the same animal. Thus, an efficient, sustainable, and humane CSF collection method is urgently needed.

In addition to assisting in CNS disease biomarker discovery, the CSF also represents an important infusion site for delivering perturbational research materials into the CNS. By bypassing the blood-brain barrier, plasmids, adenoviruses, proteins, drug molecules and cells injected into the CSF can reach the choroid plexus cell layer, brain parenchyma and spinal cord tissues at high efficiency^18–22^. These methods thus have wide potential for investigations involving the tracing and monitoring of the functional status of the CNS, establishing novel CNS disease models, and expanding our capabilities to test out therapeutics in preclinical models. Nonetheless, CSF injection in rodents remains a technically challenging task. Current CSF infusion methods are either easy to be applied on rats but not mice that have smaller body sizes while the methods for mice CSF access typically require complex surgical operations, or rely on mechanical pumps to control the rates of infusion towards the lateral ventricle or intrathecal space^18,19,23^. Considering the vast number of transgenic mouse models and their wide utilities in modeling CNS diseases, we envision that simplified CSF injection methods in the mice would significantly accelerate the development of the neuroscience field.

### Development of the protocol

A key bottleneck in the existing CSF collection and infusion methods is the requirement of surgical opening of the mouse head/neck skin to help the operator visualize the liquid chamber in the CNS and prevent over-puncturing of solid tissues by metal needles or glass capillaries. To overcome this issue, we introduce a set of bent metal needles that are specifically designed to meet the required distances between the skin and foramen magnum near the back of the neck of mice and rats. This tool constitutes the major feature in our protocol. Using this bent needle, our method does not rely on the operator’s skill to discern the changes in the resistance during the needle penetration into the CSF liquid space and transition towards solid tissues. By eliminating the necessity in visualizing the CSF liquid chamber by the operator directly, our approach bypasses the surgical opening of the rodent’s head/neck skin, which is laborious, invasive, and potentially detrimental.

Briefly, in our method, we position the head of an anesthetized animal vertically downward to expose the foramen magnum location to the operator. After identifying the foramen magnum, a bent needle is punctuated into the skin close to the edge of the foramen magnum, so the bent needle could reach to cerebellomedullary cistern. The pre-designed length of the bent needle will ensure the needle tip does not puncture and injure the brain parenchyma. Then CSF is drawn via a syringe connected to a soft tubing.

With sufficient practice, the total time required for collecting CSF from each mouse can be as short as 2-5 minutes, which significantly simplifies the procedure and reduces distress of the animal. Using this method, we frequently obtain 5-15 μL of CSF from an adult mouse and 70-120 μL from an adult rat, the volumes of which are comparable with prior terminal methods that require animal sacrifices. Through this approach, the collected CSF samples are compatible with various downstream analyses including biochemical assays and metabolomic, lipidomic and proteomic analyses.

Inspired by the minimum invasiveness and high postoperative animal survival rates of our CSF collection approach, we attempted to practice CSF injection via the foramen magnum using our bent needle tools. In this part of the method, we used a 1-cm long soft infusion tube as a connector to link a polytetrafluoroethylene (PTFE) tube to a 30G syringe bent needle. Once the connection is tightly contacted, we secure a pipette with a tip to the bent needle set, then inject 2-4 μL of desired solution such as 1% Evans blue dye into the mouse CSF. By dissecting mice at different time points after injection, we confirmed that the Evans blue dye (1% w/v) was successfully injected into the mouse CSF circulation. Notably, the ease of operation provided by the bent needle tool helped us to shorten the full injection procedure to 3-5 minutes, a dramatic improvement over prior methods. Moreover, the reduced invasiveness of the protocol allowed the animals to return to normal activity within 5-10 minutes, while none of the animals required euthanization. We envision our easy-to-practice and cost-effective method will be broadly useful for accessing the CSF in experimental animal models.

### Advantages and Limitations

#### Advantages

Key to the success of our protocol is that we replace commonly used glass capillaries with a uniquely bended metal needle made from commonly used syringes in animal experiments. Empirically, we recommend bending the needle at 90-120° with the tip length being 2-3 mm or 4-5 mm to fit with the size of an average mouse or rat respectively, to avoid accidental injury to the mice, and to reduce chances of blood contamination. Compared with prior methods, our protocol does not rely on the operator’ feeling to discern the changes in resistance during the puncturing of the micromanipulator and capillary system^15^, which is not only difficult to maneuver, but also could result in high rates of tissue injury. Of note, given the ease and accuracy provided by the handling of a bent needle, our method does not require the laborious surgical step of opening the skin and muscle covering the mouse back neck. With practice, we can accurately identify the location for needle puncturing with a near 100% success rate. With the sharp edge and thin wall of syringe needles, our tool minimizes blunt damage of the tissues surrounding the surgical area often introduced by glass capillaries. Given the simplified surgical procedure, our protocol is compatible with inhaled anesthesia, which allows the animal to recover rapidly within minutes. Together, this approach simplifies postoperative care, significantly improves the animal survival rate, and minimizes experimental cost.

Our minimally invasive CSF collection method also enables longitudinal studies that require repetitive sampling of CSF from the same individual animal. Although in our experiment, we serially collected CSF from mice in 4-day intervals and rats in 2-day intervals, which are still significantly longer than the time required for rodent CSF to be replenished^13,24^. Therefore, we expect that a shorter interval collection of CSF remains feasible. The ability to perform serial CSF collection is especially important for time-course-dependent drug efficacy investigations in animal models of CNS diseases. With simple adaptations, we believe our method can be easily applied to other laboratory animals, including hamsters, naked mole rats, and marmosets. We envision that our CSF collection method will allow new experimental design in studies involving animal models.

In addition to the collection method, our CSF injection method also greatly improves the efficiency and reduces the operative difficulty index by using our custom-made tools to inject aqueous samples into CSF via the foramen magnum. Based on the tools utilized in CSF collection, we improved the infusion tools to precisely control the injection amount by switching the syringe to a laboratory pipette connected to a PTFE capillary tube (d=0.7 mm). Furthermore, the simplified injection procedure is compatible with the inhaled anesthesia method to improve the recovery of postoperative animals. By accessing the CSF efficiently for deliveries of adenoviruses, plasmids, proteins and peptides, drugs and cells, our method will likely empower the establishment of CNS disease models in rodents more effectively.

#### Limitations

One limitation of our protocol is that the operator needs to be able to precisely locate the foramen magnum. Detailed instructions on how to identify the needle insertion site is demonstrated in our **Supplementary Videos 1 and 2**.

## Materials

### Animal

All animal experiments and procedures were conducted in accordance with institutional guidelines. All animal studies were approved by the Institutional Animal Care and Use Committee (IACUC) of Westlake University, Hangzhou, China.

- C3H/HeJ mice aged 8-9 months
- C57BL/6 mice aged 8-10 weeks
- Sprague–Dawley (SD) rats aged 7-8 months

#### Reagents

- Anesthesia: isoflurane (RWD, 3%, R510-22)
- 75% ethanol (vol/vol)
- Erythromycin eye ointment (Shuangji Beijing, 0.5%, H11021270)
- Evans blue powder (Coolaber, CE5171)

#### Equipments and Tools

- Heating pads (KEWBASIS Nanjing, KW-XMT7100)
- Animal anesthesia Vaporizer (RWD, R583S)
- Surgical table (custom-made)
- Insulin syringe (BBRAUN Omnican 40, 30G x 5/16”, 0.3 mm x 8 mm, 21B22CB)
- 1-mL disposable syringe sterile (Jiangsu Zhiyu Medical Instrument Co., LTD, 1mL, 0.45mm x 15mm)
- Sterile cotton swab (Winner, 10 cm, A1)
- Animal shaving device (Codos, CP-5200)
- PTFE capillary tube (PURESHI, Shanghai, 0.7*1.7mm)
- Microliter pipette (Eppendorf, 10μL)

### Procedure

#### Part I. CSF collection in mice and rats

##### Set up CSF collection equipment and tools. Timing: 1-2 mins

1. Connect the inhaled gas anesthesia equipment to the custom-made table and sanitize the table with 75% ethanol. If the designated table is not available, connect the inhaled gas anesthesia equipment tightly to the edge of an operation surface and secure the inhaler with tapes. (**Fig. 1a**)
2. Remove a needle from a 30G insulin syringe and hold the tip of the needle approximately 2-3 mm for mice and 4-5 mm for rats with a metal forcep. Then, using a forcep, bend the tip to a 90-120° angle. (**Fig. 1b**)
3. Cut the infusion needle from the intravenous infusion set, keeping the Luer connector. Then, connect the cutted thin tubing to the bent 30G insulin syringe needle. (**Fig. 1c**)
4. Secure a 1-mL syringe without needle (#1) to a portable holder and connect the syringe to the Luer connector of the tubing containing a bent needle. (**Fig. 1d**)
5. In addition to the above tools, prepare an extra 1-mL syringe without needle (#2) and a microcentrifuge tube for CSF collection.

**Figure 1.**
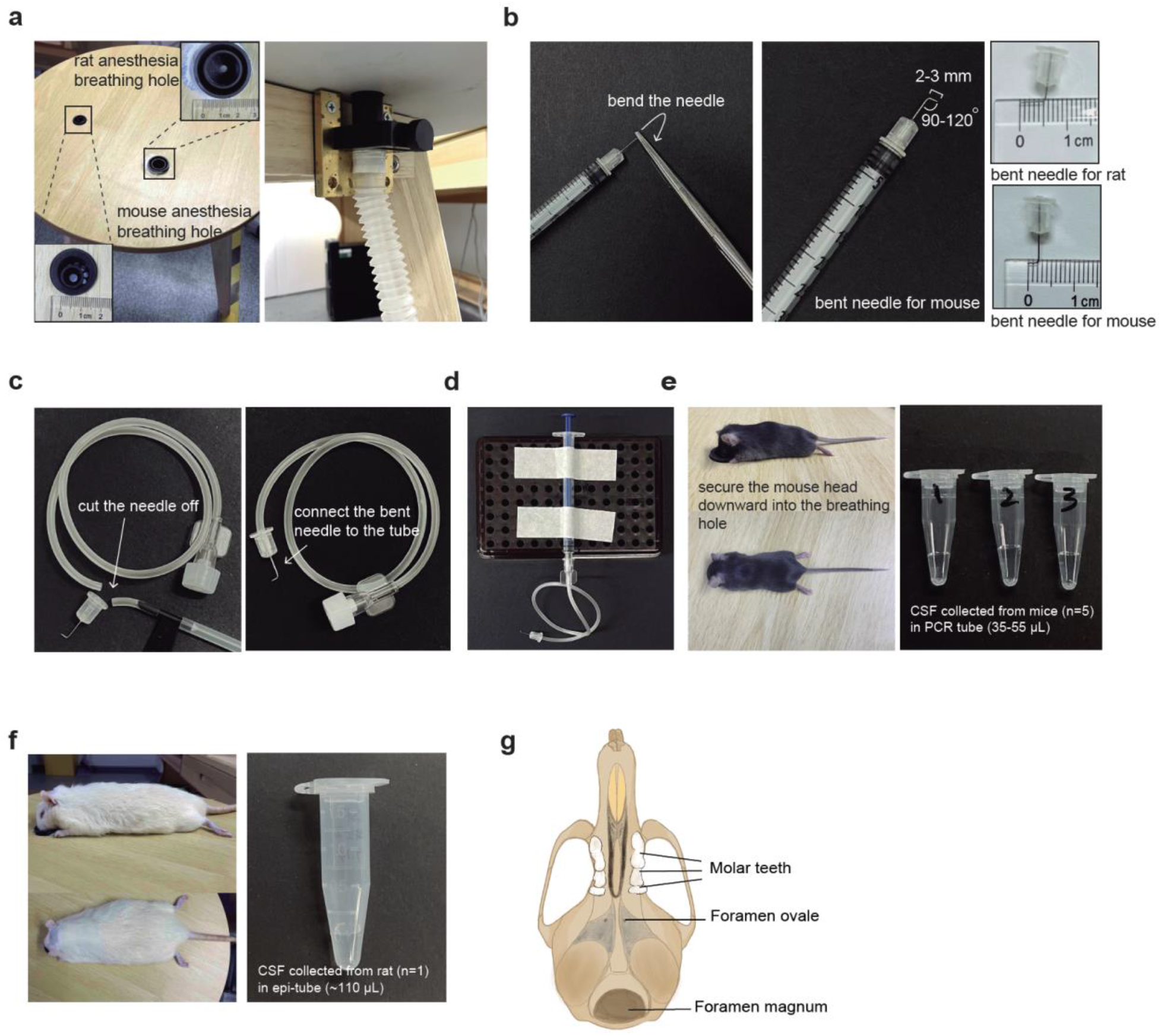
Graphical scheme and pictures describing the cerebrospinal fluid (CSF) collection procedure in mice and rats. **a.** Custom-made surgical table containing two breathable holes of different sizes for hosting mice and rats, respectively. The two holes are connected to the anesthesia machine with gas tubes. **b.** Manufacturing process of the bent needle by using a forcep to hold the needle with a certain distance and then applying force to bend. We recommend bending the needle at an angle of 90-120°. The tip length is 2-3 mm and 4-5 mm for mice and rats, respectively. **c.** Manufacturing process of surgical needle set by cutting off the needle from intravenous infusion needle, then connecting the bent needle to the soft tube. **d.** An overview of surgical tools for CSF collection by connecting custom-made surgical needle sets to a needleless 1 mL syringe. **e.** Image showing the head position of an experimental mouse under anesthesia and during the CSF collection surgery. On the right, a photo showing the CSF liquid collected from mice. Each tube contains the CSF merged from five mice, with the estimated liquid volumes (μL) indicated. **f.** Image showing the head position of an experimental rat under anesthesia and during the CSF collection surgery. On the right, a photo showing the CSF liquid collected from one rat with the estimated liquid volume indicated. **g.** Graphical scheme showing the basal view of a mouse skull, the foramen magnum is where the bent needle tip will be inserted for CSF collection in our method.

#### Mouse anesthesia and fixation. Timing 3-5 mins

##### CRITICAL See Supplementary Video 1 and 2 for specific operations

6. Anesthetize the mouse or rat by placing the animal into the induction chamber. Ensure that the animal is fully anesthetized by toe pinching reflex method.
7. Take the mouse or rat out from the chamber and record the weight of the animal using a scale.
8. Secure the head of the animal onto the custom-made surgical table. Ensure that the nose and the mouth of the animal is immediately placed into the facemasks and nosecones.
9. To avoid ocular damage during the surgery, apply erythromycin eye ointment to the eyes (optional).
10. Place the rodent’s head down vertically, so that the foramen magnum can be easily located. (**Fig. 1e,f,g**)
11. Once the location of foramen magnum is confirmed, sanitize the rodent’s head with 75% ethanol.
12. To avoid potential contamination during CSF collection, hair removal is recommended for both mice and rats.

#### CSF collection from anesthetized mouse. Timing: 1-2 mins

##### CRITICAL See Supplementary Video 1 for specific operation

13. Use one hand to hold the mouse’s head in position and use the other hand to puncture the bent needle vertically into the skin at the back of the foramen magnum. The bent needle should always be stably punctured into the skin. We recommend holding the mouse head gently and vertically downward to avoid unnecessary injury or mortality.
14. Slowly let go the hand holding the animal and gently draw the #1 syringe. By slowly pulling the plunger of the syringe, the clear CSF should be drawn into the tubing.
15. Once the CSF flow into the tubing, use the same hand to detach the tubing from the #1 syringe, and then gently remove the bent needle from the foramen magnum using the other hand.
16. Transfer the bent needle into a collection tube and attach the Luer connector of the tubing to the #2 syringe. Push the plunger and allow the CSF to flow into the microcentrifuge tube. For a one-time collection, approximately 5-15 μL and 70-120 μL of CSF can be obtained from each mouse and rat, respectively. (**Fig. 1e,f**)
17. The samples can be used for analysis immediately or stored in −80 °C freezers or in liquid nitrogen for future analysis.

#### CSF collection from rats. Timing 2-3 mins

##### CRITICAL See Supplementary Video 2 for specific operation

The procedure for rats is largely similar to mice, with the following minor differences:

18. The inhaled gas anesthesia breathable hole for rats is larger than that of the mice. Therefore, we recommend preparing two connecting tubes with different breathing holes to the anesthesia equipment and to the custom-made surgical table (**Fig. 1a**).
19. Before the operation, removing the hair covering the foramen magnum of the rat is recommended.

##### Postoperative care of the animal. Timing 5-10 mins

(The health recording period is not calculated into the surgical time)

20. After extracting CSF from the anesthetized animal, place the animal on a 37 *°C* heating pad. Typically, the animal wakes up within 1-2 mins and moves freely in a short period of time.
21. Transfer the animal into a new cage and monitor the animal for an additional 5-10 mins before placing it back to the nursing cage. We recommend monitoring the health condition and behavior of the animal on a daily basis for 7 days. When anorexia, weakness, weight loss, or infection of body organs in the animal occurs, the animal should be cared for according to animal welfare guidelines or euthanized.

#### Part II. CSF injection in mice

##### Evans Blue dye solution (1% w/v) preparation. Timing: 1-2 mins

1. Weight 0.1 g of evans blue powder and dissolve in 10 mL of sterile saline.

##### Set up of CSF injection equipment and tools. Timing: 1-2 mins

2. Connect the inhaled gas anesthesia equipment to the custom-made table as described in the CSF collection section.
3. Prepare the bent needle as described in the CSF collection section.
4. Cut approximately 1-cm long tube from the intravenous infusion set as a connector tube. Insert the PTFE tube into the connector tube about 5mm deep. Once the insertion is securely done, attach the bent 30G insulin syringe needle to the other end of the connector tube. (**Fig. 4a**)
5. Insert a 10-μL pipette tip to the other end of PTFE capillary tube and tightly secure the pipette tip to the pipette. (**Fig. 4b**)
6. Immerse the bent needle of the connected tubing set into the 1% Evans blue dye solution (w/v) and aspirate 3 μL of 1% Evans blue dye solution (w/v) into the PTFE capillary tube for injection.

##### Mouse anesthesia and fixation. Timing 3-5 mins

7. Detailed information please refer to step 6-12 of the CSF collection part.

##### CSF injection with anesthetized mouse. Timing: 1-2 mins

8. Use one hand to hold the mouse’s head in position and use the other hand to puncture the bent needle vertically into the skin at the back of the foramen magnum. Normally, when the bent needle punctures through foramen magnum and inserts to cerebellomedullary cistern, the fluid level in PTFE capillary tube rises.
9. Slowly let go the hand holding the animal and gently push the pipette. By slowly pushing the pipette, the Evans blue dye solution (1% w/v) should be injected into the CSF circulation.

##### Postoperative care of the animal. Timing 5-10 mins

10. After injecting 1% (w/v) Evans blue solution into CSF from the anesthetized animal, place the animal on a 37 *°C* heating pad. Typically, the animal wakes up within 5-10 mins and moves freely in a short period of time.
11. According to experimental needs, post-operative mice can be either dissected or transferred back to the nursing cage. For nursing, we recommend monitoring the health condition and behavior of the animal on a daily basis for at least 7 days. When anorexia, weakness, weight loss, or infection of body organs in the animal occurs, the animal should be cared for according to animal welfare guidelines or euthanized.

##### Troubleshooting (Part I. CSF collection)

| Step | Problem | Possible reason | Solution |
| --- | --- | --- | --- |
| 10 | Difficulty in identifying the location of foramen magnum | The position of the animal's head is not vertically down | Hold the neck of the animal and gently push the head forward, while using the other hand to feel the location of the foramen magnum, until finding a favorable position to operate the surgery. |
| 14 | The punctured needle moves once removing hand away from holding the animal | The hand holding the punctured bent needle is not stable enough | Apply a slight force downward to animal's head by the hand holding the bent needle, make sure that the needle is stably punctured into skin without moving |
| 14 | Blood goes into the tube | Needle punctures a blood vessel | Puncture the needle vertically and do not move the needle during the process |
| 15 | CSF mixed with air bubbles is quickly drawn into the connection tube | Pull out the bent needle before removing the connection tube from syringe, so the CSF is sucked due to the negative pressure inside the tube | Remove the connection tube from syringe first, then remove the bent needle from animal |

##### Troubleshooting (Part II. CSF injection)

| Step | Problem | Possible reason | Solution |
| --- | --- | --- | --- |
| 6 | Evans blue solution may leak out when aspirate with custom-made bent needle set | The link between PTFE tube and 30G syringe needle is not tightly connected by the connector tube | With the help of a connector tube, try to insert the PTFE tube closely connected to the 30G syringe needle; or directly connect PTFE tube to the 30G syringe needle with force |
| 8 | Difficulty in observing the rise of fluid level in PTFE capillary tube | The puncturing location is not correct | Adjust the puncture location to allow the needle inserting into the skin through the edge of the foramen magnum and reaching to cerebellomedullary cistern |
| 9 | Difficulty in pushing the pipette and injecting liquid samples | The needle may have punctured into the muscle tissue instead of the circulation cavity of CSF | Same as the solution for step 8. |

##### Timing (Part I. CSF collection)

Step 1-5, set up the equipments and tools: 1-2 mins

Step 6-12, animal anesthesia and fixation: 3-5 mins

Step 13-19, mouse or rat CSF collection: 2-3 mins

Step 20, observation after the operation: 5-10 mins

Step 21, post-operative care: monitoring for 7 days

##### Timing (Part II. CSF injection)

Step 1, Evans blue dye solution (1% w/v) preparation: 1-2 mins

Step 2-6, set up CSF injection equipment and tools: 1-2 mins

Step 7, mouse anesthesia and fixation: 3-5 mins

Step 8-9, CSF injection with anesthetized mouse: 1-2 mins

Step10-11, post-operative monitoring: 5-10 mins

### Anticipated results

Collecting CSF from small animals, especially mice, has been technically challenging due to the sophisticated surgical procedures and high mortality rates. Using our bent needle method, we can consistently collect 5-15 μL of CSF from adult mice and 70-120 μL from adult rats (**Fig. 1e,f**). To ensure our method causes minimal invasion to the animal, we monitored the recovery rate and survival rate of postoperative mice. In our CSF collection experiment, mice typically recover from anesthesia within 1-5 min without notable health issues. This is a significant achievement compared with prior methods.

Encouraged by the rapid postoperative recovery of rodents, we attempted to collect CSF repetitively from the same animal using our method after sufficient recovery time. The experiment involved three CSF collections in a group of 15 adult mice at day 0, 4, 8, respectively, and all collections were successful without causing notable body weight loss, other health issues or death (**Fig. 2a,b**). The volumes of CSF extracted in later collections were comparable to those in the first collection. These mice continued to survive through another 10-days of monitoring period without showing any notable signs of distress (**Fig. 2a,b**). Similarly, rats used in CSF collection also demonstrated a rapid post-surgical recovery and a stable survival from sequential CSF extraction at day 0, 2, 4, respectively **(Fig. 2c).** These results support that our method is compatible with longitudinal studies requiring serial CSF collection from the same animal, a major advantage over prior protocols that require extensive surgical operations.

**Figure 2.**
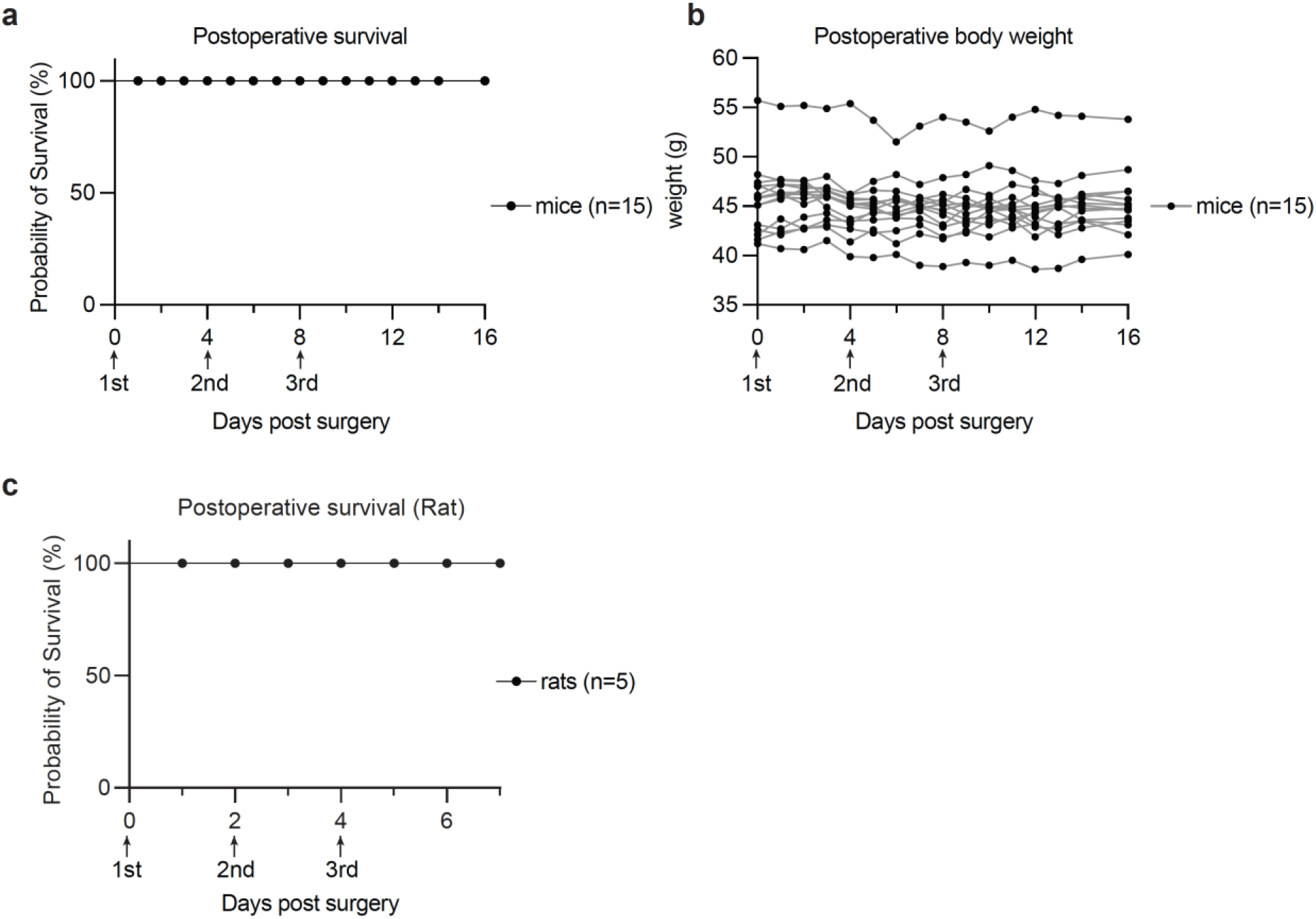
Animal survival and body weight monitoring after CSF collection. **a.** Kaplan-Meier curve showing the survival rate of postoperative mice (n=15). **b.** Curves showing the time-course monitoring of body weights in the postoperative mice used in **Figure a**. **c.** Kaplan-Meier curve showing the survival rate of postoperative rats (n=5).

To validate our technique, we performed proteomic analysis to identify previously-characterized marker molecules in the isolated CSF solutions (**Fig. 3a**). Mass spectrometry analysis detected at least 513 proteins in the pooled mouse CSF samples (**Fig. 3a, Supplementary Dataset 1**). These proteins include Apolipoprotein E (Apoe) and Apolipoprotein A-1 (Apoa1) – the predominant apolipoproteins in mammalian CSF^25^, Amyloid beta precursor protein (App)^26^, Serotransferrin (Tf)^27^, Transthyretin (Ttr)^27^, Prostaglandin D2 Synthase (Ptgds)^27^, etc (**Fig. 3a**). These molecular features indicate that we were successfully collecting the cerebrospinal fluid from the mouse.

**Figure 3.**
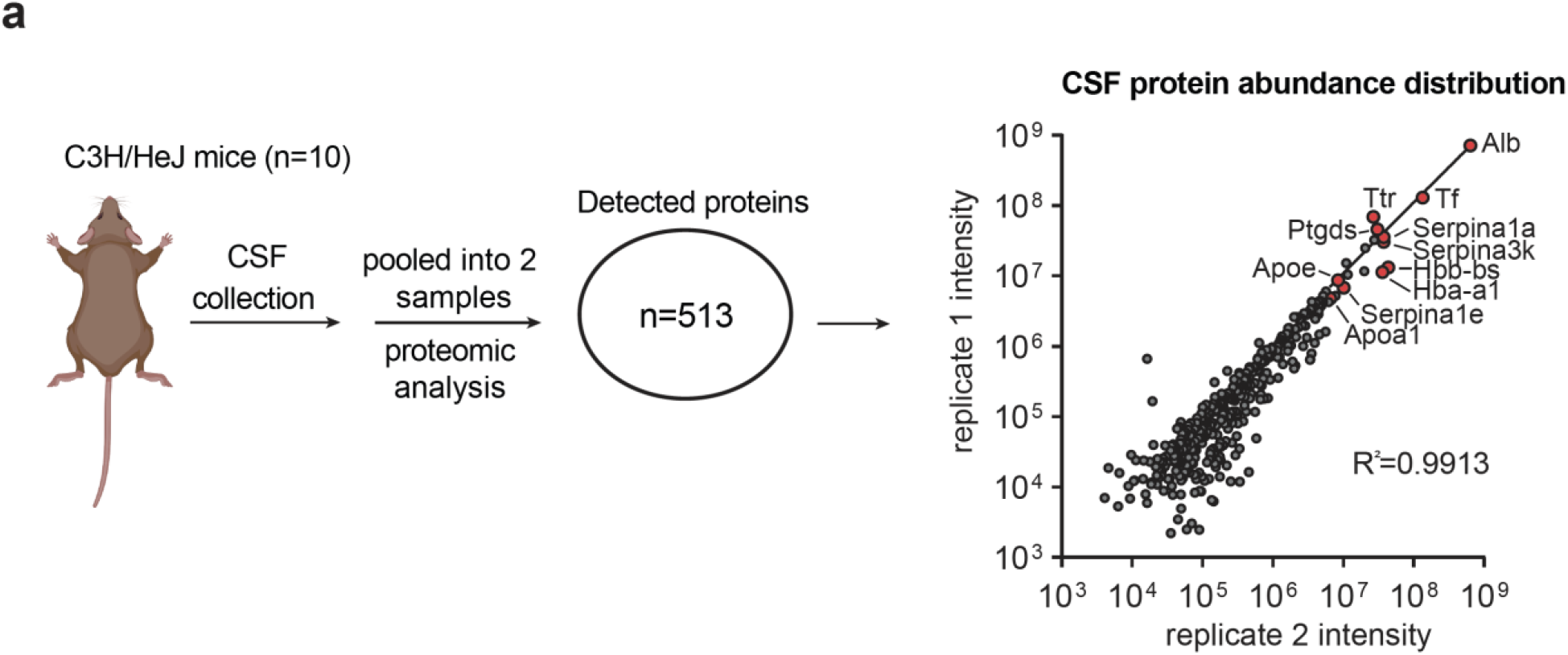
Proteomics analysis of mouse CSF collected using the present method. **a.** Graphical scheme and scatter plot describing the proteomics analysis of CSF collected from adult mice and the CSF proteome abundances from two replicate pooled samples. Each sample contains the CSF pooled from 5 mice. Highly abundant proteins detected in both samples are selectively labeled in red. Correlation analysis is performed using the Pearson correlation method.

We used the Evans blue dye, a metabolically inert, colored marker to indicate whether our CSF injection was successful (**Fig. 4c**). After dissecting the mice brain at 1, 2 or 3 hr post-injection, we found that the dye readily diffused away from the injection site at the foramen magnum and covered a major part of the brain surface, which fits with the coverage of the CSF circulation (**Fig. 4c**). This result indicates that our bent needle injection method can be used to deliver liquid reagents at the lower μL scale to the mouse CSF circulation.

**Figure 4.**
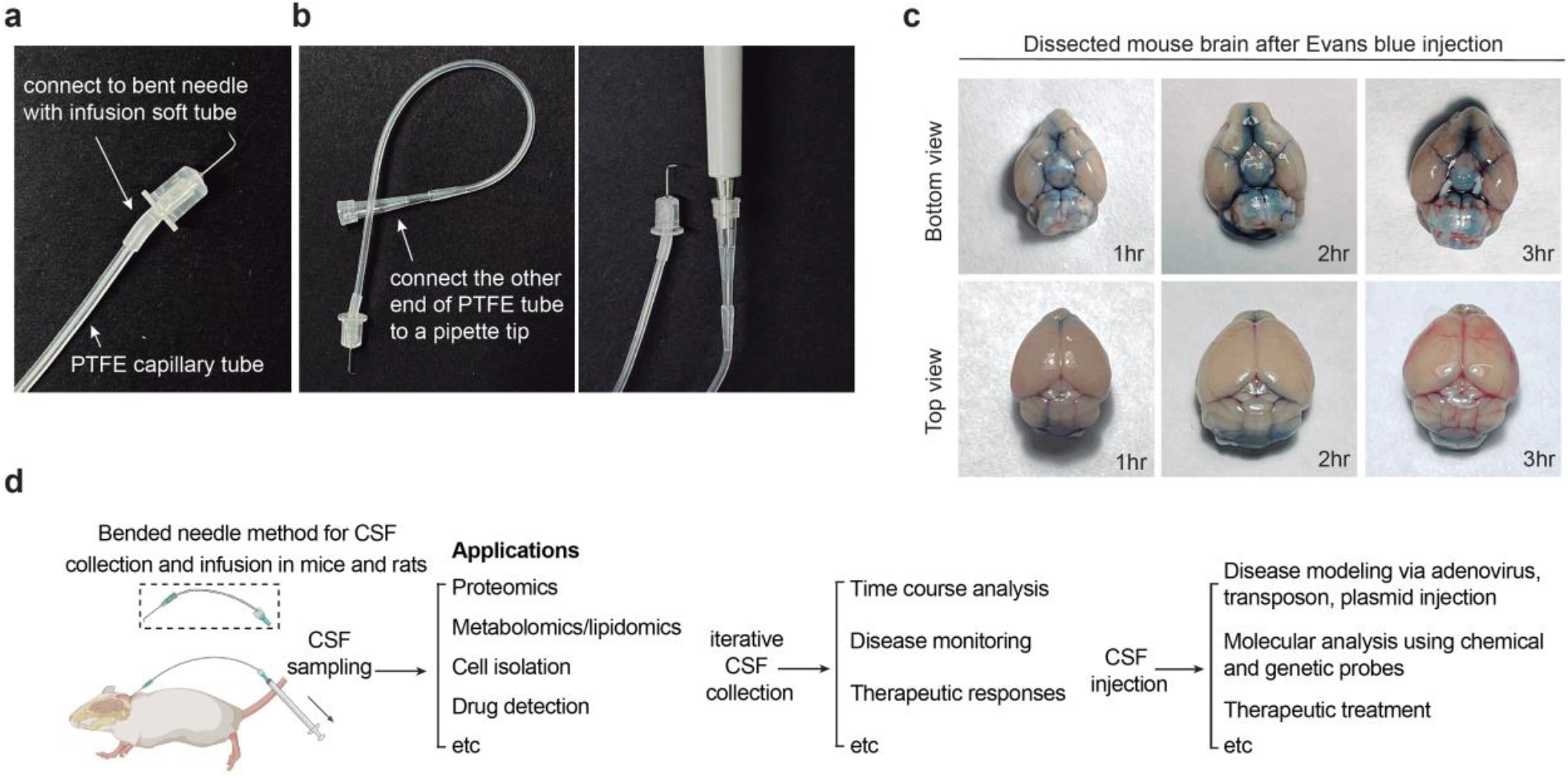
Procedures for CSF injection in mice. **a.** Graphical scheme for the injection tools by connecting the bent needle with infusion soft tube to the PTFE capillary tube. Once the bent needle end is fully made, connect the other end of the PTFE tube to a 10 μL pipette tip and a pipette. **b.** Optical imaging of the dissected mouse brains 1, 2, and 3 hr after injection of 1% Evans blue solution. The brain images are depicted at both top and bottom views. **c.** Scheme showing the proposed utilities of the developed CSF collection and infusion methods.

### Applications

By introducing the key bent needle tool described above, we envision that the minimally invasive CSF collection and infusion methods presented here will enable more efficient monitoring of the metabolic and functional states of the central nervous system in research involving small laboratory animals including mice and rats (**Fig. 4d**). This approach allows multi-omic profiling of the collected CSF including proteomics, metabolomics, lipidomics as well as cell isolation for characterization of the CSF compositions. Notably, iterative CSF collection from the same animal will enable new experimental strategies that were previously not possible or too laborious and costly, including time-course monitoring of disease progression and therapeutic responses (**Fig. 4d**). Moreover, our method is compatible with delivering various reagents into the CSF, including plasmids, chemical probes, proteins and peptides and adenoviruses, which will likely improve the technical efficiency in CNS disease modeling, tracing and monitoring of CNS functions, as well as imposing therapeutic treatments. We envision that our technique will be widely useful at the forefronts of neuroscience.

## Supplementary Videos

**Supplementary Video 1. Video illustrating the cerebrospinal fluid collection process in a mouse.**

**Supplementary Video 2. Video illustrating the cerebrospinal fluid collection process in a rat.**

## Supplementary Datasets

**Supplementary Dataset 1. Proteomic analysis dataset of mouse CSF samples.**

## Supplementary Information

### Methods for proteomic analysis of mouse CSF samples

#### Sample preparation

10 μL of each cerebrospinal fluid sample was transferred to a 1.5 mL centrifuge tube and lysed with 50 μL of lysis buffer (8 M Urea, 100 mM TEAB, pH 8.5). The lysate was centrifuged at 12,000 g for 15 min at 4 °C, and the supernatant was reduced with 10 mM DTT for 1 hr at 56 °C. The samples were subsequently alkylated with sufficient IAM for 1 hr at room temperature in the dark.

#### Trypsin digestion

The volume of each protein sample was made up to 100 μL with lysis buffer (8 M Urea, 100 mM TEAB, pH 8.5). Trypsin and 100 mM TEAB buffer were added, and the samples were mixed and digested at 37 °C for 4 hr. Then trypsin was replenished and CaCl2 was added to digest overnight. Formic acid was mixed with a digested sample, adjusted pH under 3, and centrifuged at 12,000 g for 5 min at room temperature. The supernatant was slowly loaded to the C18 desalting column, washed with a washing buffer (0.1% formic acid, 3% acetonitrile) for 3 times, then added an elution buffer (0.1% formic acid, 70% acetonitrile). The eluents of each sample were collected and lyophilized.

#### Separation of fractions

Mobile phase A (2% acetonitrile, adjusted pH to 10.0 using ammonium hydroxide) and B (98% acetonitrile, adjusted pH to 10.0 using ammonium hydroxide) were used to develop a gradient elution. The lyophilized powder was dissolved in solution A and centrifuged at 12,000 g for 10 min at room temperature. The sample was fractionated using a C18 column (Waters BEH C18, 4.6×250 mm, 5 μm) on a Rigol L3000 HPLC system; the column oven was set as 45 °C. The detail of the elution gradient was shown in Table 1. The eluates were monitored at UV 214 nm, collected for a tube per minute and combined into 10 fractions finally. All fractions were dried under vacuum, and then, reconstituted in 0.1% (v/v) formic acid (FA) in water.

**Table 1.** nanoElute Liquid chromatography elution gradient table

| Time (min) | Flow rate (mL/min) | Mobile phase A (%) | Mobile phase B (%) |
| --- | --- | --- | --- |
| 0 | 300 | 98 | 2 |
| 45 | 300 | 78 | 22 |
| 50 | 300 | 65 | 35 |
| 55 | 300 | 20 | 80 |
| 60 | 300 | 20 | 80 |
| 75 | 300 | 78 | 22 |
| 80 | 300 | 63 | 37 |
| 85 | 300 | 5 | 95 |
| 90 | 300 | 5 | 95 |

#### LC-MS/MS Analysis

Mobile phase A (100% water, 0.1% formic acid) and B solution (100% acetonitrile, 0.1% formic acid) were prepared. The lyophilized powder was dissolved in 10 μL of solution A, centrifuged at 14,000 g for 20 min at 4 °C, and 200 ng of the supernatant was injected into the Liquid chromatography-mass spectrometry system to detect. The model type of the UHPLC was nanoElute with nano-upgraded, and the analytical column was a home-made analytical column (15 cm×100 μm, 1.9 μm). The elution conditions of liquid chromatography were using a 90 min elution gradient method, which contains 300 flow rate (mL/min), 5% mobile phase A, and 95 % mobile phase B. Mass spectrometry analysis was performed on a tims TOF Pro2 mass analyzer (Bruker) with Captive Spray ion source. The spray voltage was set to 2.1 kV. The full scan range of the mass was from *m/z* 100 to 1700 and the Ramp time was 100 ms. The Lock Duty Cycle was set to 100%. The settings of PASEF were as follows: 10 MS/MS scan (a total cycle time of 1.17 sec), ionic strength threshold of 2500, scheduling target intensity of 20000. The raw data of MS detection was named as “.d”.

#### Data analysis

The resulting spectra were searched against the Uniprot mouse proteome database by the MaxQuant (Bruker) search engines. The search parameters of MaxQuant are set as follows: mass tolerance for precursor ion was 20 ppm and mass tolerance for product ion was 0.05 Da. Carbamidomethyl was specified as fixed modifications, Oxidation of methionine (M) was specified as dynamic modification, and acetylation was specified as N-Terminal modification. A maximum of 2 missed cleavage sites were allowed. In order to improve the quality of analysis results, the software PD or MaxQuant further filtered the retrieval results: Peptide Spectrum Matches (PSMs) with a credibility of more than 99%were identified as PSMs. The identified protein contains at least 1 unique peptide. The identified PSMs and protein were retained and performed with FDR no more than 1.0%.

## Acknowledgements

We thank Dr. Jingjing Bao and the Laboratory Animal Resource Center at Westlake University for providing technical support, and Novogene (Beijing) for assisting proteomic analysis of CSF samples. We thank members of the Zou lab for insightful discussions. This work was supported by Westlake Laboratory of Life Sciences and Biomedicine (for Y.Z.) and Startup funds from Westlake Education Foundation (for Y.Z.).

## Author contributions

J.R.H. and Y.Y. designed and performed the experiments. T.W.W., T-T.S., and W.L. assisted the initial method development. Y.Z. supervised the project. All authors contributed to manuscript preparation.

## Additional Information

Correspondence and requests for materials should be addressed to Y.Z.

